# Identification of novel inhibitors of DLK palmitoylation by High Content Screening

**DOI:** 10.1101/430629

**Authors:** Dale D.O. Martin, Prasad S. Kanuparthi, Sabrina M. Holland, Shaun S. Sanders, Hey-Kyeong Jeong, Marget B. Einarson, Marlene Jacobson, Gareth M. Thomas

**Affiliations:** Shriners Hospitals Pediatric Research Center, Lewis Katz School of Medicine at Temple University, 3500 N. Broad Street, Philadelphia, PA 19140.; Fox Chase Cancer Center, Philadelphia, PA; Moulder Center for Drug Discovery Research, Temple University School of Pharmacy; Department of Anatomy and Cell Biology, Lewis Katz School of Medicine at Temple University

## Abstract

After axonal insult and injury, Dual leucine-zipper kinase (DLK) conveys retrograde pro-degenerative signals to neuronal cell bodies *via* its downstream target c-Jun N-terminal kinase (JNK). We recently reported that such signals critically require modification of DLK by the fatty acid palmitate, *via* a process called palmitoylation. Compounds that inhibit DLK palmitoylation could thus reduce neurodegeneration, but identifying such inhibitors requires a suitable assay. Here we report that DLK subcellular localization in non-neuronal cells is highly palmitoylation-dependent and can be used as a proxy readout to identify inhibitors of DLK palmitoylation by High Content Screening (HCS). We exploited this highly specific localization of DLK-GFP as the basis for a screen of the Prestwick Compound Library™. We found that ketoconazole, a Prestwick Library compound that most dramatically affected DLK subcellular localization in our primary screen, inhibited DLK palmitoylation in a dose-dependent manner in follow-up biochemical assays. Moeroever, ketoconazole significantly blunted phosphorylation of c-Jun in primary sensory neurons subjected to Trophic Deprivation, a well known model of DLK-dependent pro-degenerative signaling. These findings suggest that our HCS platform is capable of identifying novel inhibitors of DLK palmitoylation and signalling that may have considerable therapeutic potential.

## Introduction

In both chronic neuropathological conditions and following acute injury, Dual Leucine-zipper Kinase (DLK) signals *via* its downstream target c-Jun N-terminal Kinase (JNK) to activate pro-degenerative transcription and subsequent neuronal death [1–7]. Genetic knockout of DLK confers striking neuroprotection in several models of neurodegeneration, spurring great interest in targeting DLK therapeutically as a neuroprotective strategy [1, 2, 5, 7]. Indeed, inhibitors of DLK’s kinase activity have shown therapeutic promise in multiple animal models of disease [1, 8–10]. Unfortunately, though, the most promising DLK inhibitors reported thus far also inhibit additional kinases [8], which may limit the potential of this therapeutic approach.

An alternative or complementary strategy that holds considerable promise would be to target DLK-specific regulatory features. Our studies of DLK-specific regulation led to our recent finding that DLK undergoes palmitoylation [11], the reversible covalent addition of a saturated fatty acid, typically palmitate [12–14]. Palmitoylation is best known to control protein subcellular localization and we found that palmitoylation targets DLK to specific axonal vesicles in primary sensory neurons [11]. ‘Hitch-hiking’ on these vesicles may allow DLK to convey retrograde signals from damaged axons to neuronal cell bodies [11]. Interestingly, though, palmitoylation plays an unexpected additional role, because it is also critical for DLK to phosphorylate and activate ‘downstream’ JNK pathway kinases [11]. Consistent with the importance of palmitoylation for DLK-JNK signaling, genetically mutating DLK’s palmitoylation site prevented JNK phosphorylation in non-neuronal cells, and blocked JNK-dependent responses to axonal injury in cultured neurons [11]. These findings suggested to us that compounds that prevent DLK palmitoylation might be as neuroprotective as inhibitors of DLK’s kinase activity. However, pursuing this therapeutic strategy would require development of an effective screening method to identify such compounds.

Here we report that in non-neuronal cells, DLK localization is also highly palmitoylation-dependent. This localization can be used as a proxy for DLK palmitoylation that is compatible with a High Content Screening (HCS) approach. We optimized our screen to identify and eliminate compounds that broadly affect protein transcription, translation and/or stability and to eliminate likely cytotoxic compounds. Using these optimized conditions we screened a library of FDA-approved compounds and identified several that specifically affect DLK localization. Ketoconazole, an antifungal agent that most dramatically affected DLK localization in our primary screen, also inhibited DLK palmitoylation in follow-up biochemical assays and reduced DLK-dependent signaling in primary neurons. Our screening assay thus has the potential to identify novel modulators of DLK palmitoylation, which may have considerable therapeutic potential.

## Results

### DLK subcellular localization is highly palmitoylation-dependent in HEK293T cells

In primary sensory neurons, DLK localizes to axonal vesicles [11]. This discrete localization is prevented by a pharmacological inhibitor of protein palmitoylation (the compound 2-Bromopalmitate (2BP; [15])) or by point mutation of DLK’s palmitoylation site, Cys-127 [11]. Subcellular localization changes of this type are often used as readouts in High Content Screening (HCS) [16, 17], an approach that might therefore be well suited to identify compounds that inhibit DLK palmitoylation. However, because a non-neuronal cell line might be more amenable to HCS approaches than primary neurons, we assessed whether DLK localization is also palmitoylation-dependent in HEK293T cells. We found that transfected wild type GFP-tagged DLK (wtDLK-GFP) in HEK293T cells localizes to intracellular membranes that colocalize with the Golgi marker GM130 (Figure 1A). wtDLK’s Golgi localization in HEK293T cells may be because the axonal vesicle population is not present in this cell line and/or because many mammalian palmitoyl acyltranserases (PATs, which catalyze palmitoylation) localize to the Golgi in these cells [18]. Importantly, though, this localization was again highly palmitoylation-dependent, because both 2BP treatment and C127S mutation shifted DLK localization from Golgi-associated to diffuse (Figure 1B).

**Figure 1.**
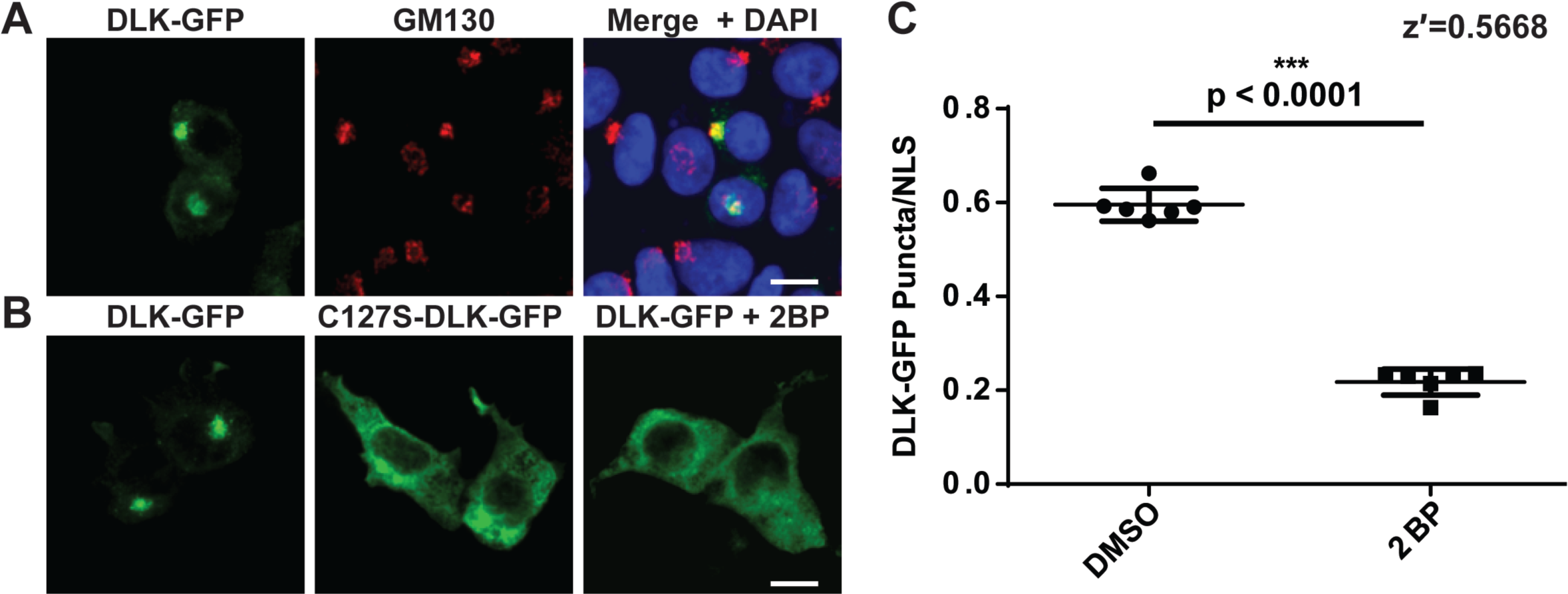
Palmitoylation-dependent localization of DLK-GFP to the Golgi apparatus in HEK293T cells. **A**) HEK293T cells were transfected to express wild type DLK-GFP and subsequently fixed. DLK-GFP and the Golgi marker GM130 were detected with specific antibodies and nuclei were detected using the DNA dye DAPI. **B**) HEK293T cells were transfected as in *A* to express either wild type DLK-GFP (DLK-GFP) or a DLK palmitoylation site mutant (C127S-DLK-GFP). C127S mutation, or treatment with the palmitoylation inhibitor 2BP, diffuses the Golgi-associated clusters of DLK-GFP. **C**) HEK293T cells were seeded into 12 wells of a 96-well plate and transfected with DLK-GFP and NLS-mCh cDNA and then treated with 2BP (20 μM in DMSO) or 0.1 % (v/v) DMSO vehicle (6 wells per condition). Cells were fixed and imaged using an ImageXpress High Content Imaging system to detect GFP signal. Assay quality was determined by calculating the z-prime (z′) for 6 determinations for each of the indicated conditions (z′=S/R, S = [(Mean of Vehicle treated − 3xSD)-(Mean of 2BP − 3xSD)], R = Vehicle Mean − 2BP mean).

### Palmitoylation-dependent control of DLK localization is a robust, HCS-compatible readout

Given that 2BP treatment and C127S mutation completely eliminate DLK palmitoylation in biochemical assays [11], Golgi localization of DLK can thus serve as an effective proxy for DLK palmitoylation that may be HCS-compatible. We therefore sought to quantify the DLK localization change using HCS software. Images of DLK-GFP fluorescence were acquired using an ImageXpress High Content Analyzer and each was then thresholded to an identical value. Under these conditions, numerous Golgi-associated punctate structures (henceforth ‘puncta’) of wtDLK-GFP could be observed and quantified. These puncta were essentially absent in the 2BP-treated condition (Figure 1C). The DLK-GFP localization change is therefore an HCS-compatible proxy readout of DLK palmitoylation.

We next assessed the robustness of our assay by calculating the z-prime (z′), a statistical measurement commonly used to evaluate and validate HCS assays [19]. Typically, a z′ value of ≥0.5 is deemed an excellent assay. We therefore seeded HEK293T cells in 96 well plates, prior to transfection with wtDLK-GFP and subsequent treatment with 2BP (positive control) or vehicle. After fixation and subsequent ImageXpress analysis, we calculated a z′ value of 0.57 (Figure 1C), indicative of a highly robust, HCS-compatible assay. Additional metrics of DLK’s subcellular distribution, including the area per field of view occupied by punctate structures and the average intensity of these puncta, also showed a high degree of palmitoylation-dependence (Supplementary Fig 1).

### An optimized high-throughput imaging screen for DLK palmitoylation

Our goal in establishing the HCS assay was to identify compounds that reduce DLK puncta number because they reduce DLK palmitoylation. However, we realized that DLK puncta numbers might also be reduced by compounds that impaired DLK transcription or translation, or by cytotoxicity. To facilitate detection of ‘false positive’ compounds that broadly affect these processes, we therefore adapted our assay to incorporate a cotransfected cDNA that codes for a nuclear localization signal (NLS) fused to the red fluorescent reporter mCherry (mCherry-NLS) [20], expressed downstream of the same CMV promoter used in the DLK-GFP cDNA. We also included the nuclear marker DAPI to quantify healthy nuclei per well. Reduced DAPI counts and/or nuclear fragmentation (which is detectable by ImageXpress software) can serve as an additional indicator of potentially cytotoxic compounds. Importantly, 2BP affected neither mCherry-NLS nor DAPI counts at the concentration used (Figure 2A).

**Figure 2.**
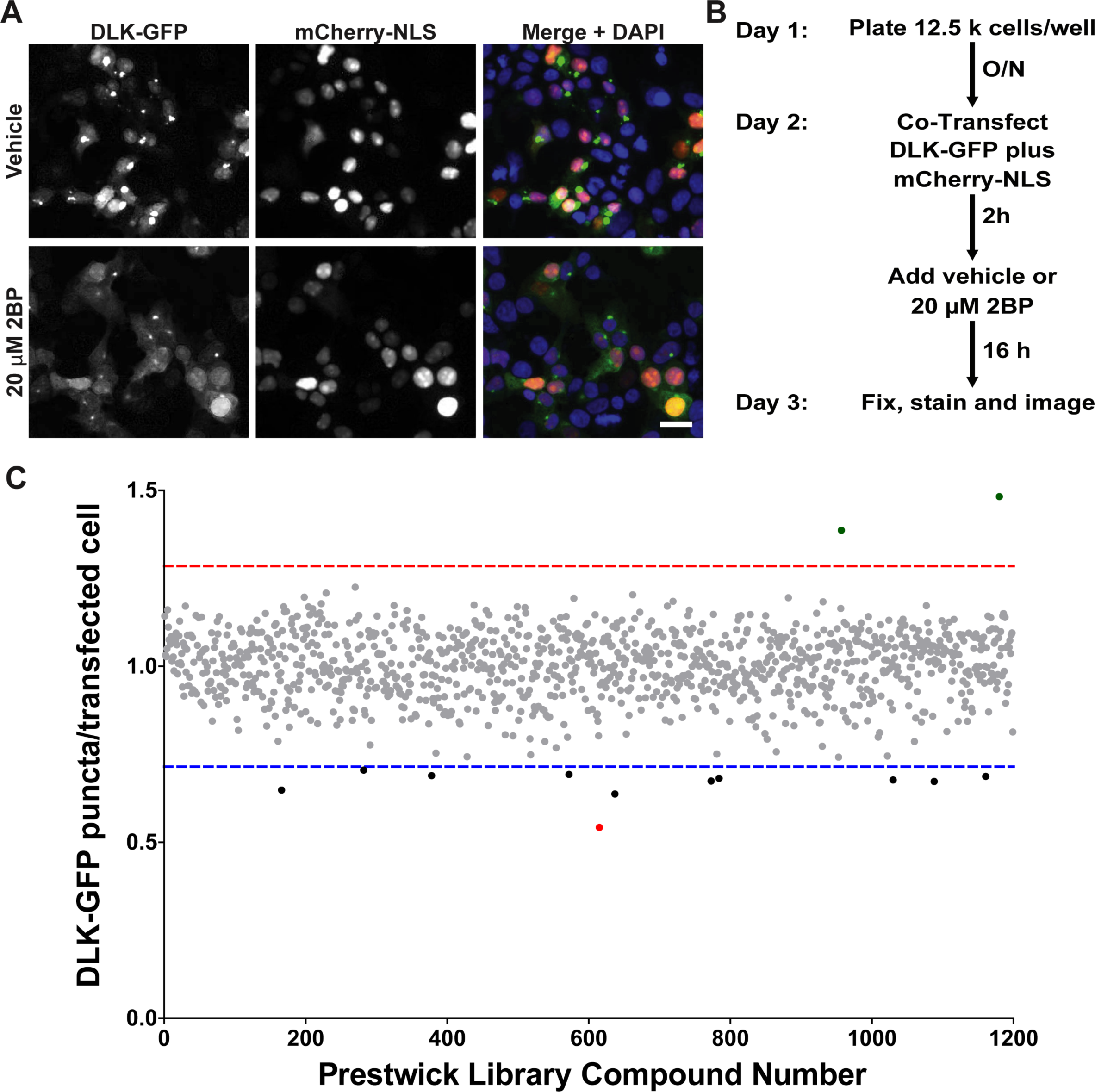
A High Content Imaging screen identifies ketoconazole as the most potent compound to inhibit DLK-GFP puncta formation. **A**) HEK293T cells cotransfected with DLK-GFP plus mCherry-NLS were treated with 2BP or vehicle and fixed to detect GFP, mCherry and the nuclear marker DAPI. 2BP reduces DLK-GFP puncta without affecting mCherry-NLS expression or DAPI signal. Scale bar: 25 μm. **B**) Design of the high-throughput screen for compounds that inhibit DLK-GFP punctate localization **C**) The effect of 1200 compounds from the Prestwick Chemical Library™ on DLK puncta per transfected cell (mean of 2 determinations per compound) was calculated using ImageXPress Image Analysis ‘TransFluor’ and Multi-Wavelength Scoring (MWS) modules. Compounds that decreased the number of transfected cells (from mCherry-NLS count) or the total number of cells (from DAPI count) by greater than 30%, relative to the mean of vehicle treated controls, are not plotted due to likely cytotoxicity or non-specific effects. Red and blue dotted lines indicate 3 standard deviations (3SD) above and below the mean of all determinations, respectively. Compounds that decreased DLK puncta per transfected cell below this 3SD cut-off were considered ‘Hits’. The most potent ‘hit’, ketoconazole, is highlighted in red.

### A Prestwick Chemical Library™ screen reveals that the compound Ketoconazole reduces DLK punctate localization

Having validated our HCS approach with the positive control ‘tool compound’ 2BP, we sought to expand our assay to perform an initial library screen. The Prestwick Chemical Library™ consists of over 1200 compounds that have been approved by the FDA, EMA, or other agencies for use in humans [21]. The library was prepared by medicinal chemists and pharmacists to ensure high chemical diversity and known bioavailability in humans, thereby increasing the likelihood of identifying “high quality” hits. Another advantage of the Prestwick library is that positive hits have the potential to be used immediately in downstream analyses and studies.

Using our optimized conditions we therefore expanded our assay to screen the 1200 Prestwick library compounds (Figure 2B). Compounds that reduced either mCherry-NLS or DAPI signals by more than 30%, relative to the mean of vehicle-treated controls, were excluded from further analysis due to likely cytotoxicity or effects on transcription/translation/protein stability. The effect of all remaining compounds on DLK puncta, relative to the total mCherry-NLS count (“DLK puncta per transfected cell”), was then quantified (Figure 2C). Eleven compounds reduced DLK puncta per transfected cell by > 3X SD, relative to the mean value for all compounds (Table 1). The antifungal compound ketoconazole had the greatest effect, reducing DLK puncta per transfected cell by 45.8 ± 0.5%, and was selected for follow-up studies.

**Table 1.** Compounds identified in the Prestwick Compound Library that reduced DLK-GFP puncta per transfected cell (“Puncta/NLS”) by >3xSD, relative to the mean of all compounds tested.

| Rank | Drug | Puncta/NLS<br>Read 1 | Puncta/NLS<br>Read 2 | Average<br>Puncta/NLS | Notes |
| --- | --- | --- | --- | --- | --- |
| 1 | Ketoconazole | 0.537 | 0.547 | 0.542 | Antifungal. Inhibits 17α-hydroxylase and 17,20-lyase, which are involved steroid synthesis, including testosterone precursors |
| 2 | Lidoflazine | 0.687 | 0.588 | 0.637 | Ca+2 channel blocker. Coronary vasodilator. |
| 3 | Bromocryptine mesylate | 0.655 | 0.642 | 0.649 | Also known as Parlodel. Dopamine agonist. |
| 4 | Thioguanosine | 0.694 | 0.652 | 0.673 | Chemotherapeutic |
| 5 | Nefazodone HCl | 0.688 | 0.661 | 0.674 | 5-HT2A receptor antagonist |
| 6 | Sulconazole nitrate | 0.662 | 0.692 | 0.677 | Also known as Exelderm. Antifungal |
| 7 | Nifedipine | 0.671 | 0.693 | 0.682 | Also known as Adalat. Ca+2 channel blocker. |
| 8 | Tyloxapol | 0.679 | 0.696 | 0.687 | Non-ionic polymer. Component of the expectorant Tacholiquin. |
| 9 | Disulfiram | 0.653 | 0.727 | 0.690 | Also known as Antabuse and Antabus. Inhibits acetaldehyde dehydrogenase. |
| 10 | Imatinib | 0.665 | 0.721 | 0.693 | Also known as Gleevec. Inhibits bcr-abl for treatment of chronic myelogenous leukemia |
| 11 | Clomiphene citrate (ZE) | 0.687 | 0.723 | 0.705 | Selective estrogen receptor modulator (SERM). |

### Ketoconazole inhibits DLK puncta formation and palmitoylation

We first assessed the dose-dependence of ketoconazole’s effect on DLK localization using a re-purchased stock of the compound. At the concentration used in the initial screen (10 μM) Ketoconazole again greatly reduced the number of DLK-GFP puncta/transfected cell (Fig 3A). Ketoconazole’s effect on DLK-GFP puncta/transfected cell was clearly dose-dependent, first reaching statistical significance at 2.5 μM (Fig. 3A). At concentrations >10 μM, ketoconazole more markedly reduced the number of DLK-GFP puncta/transfected cell, but also clearly affected the number of transfected cells. However, these findings suggested that a clear window exists within which ketoconazole reduces DLK-GFP puncta/transfected cell without affecting overall transcription/translation/protein stability.

**Figure 3.**
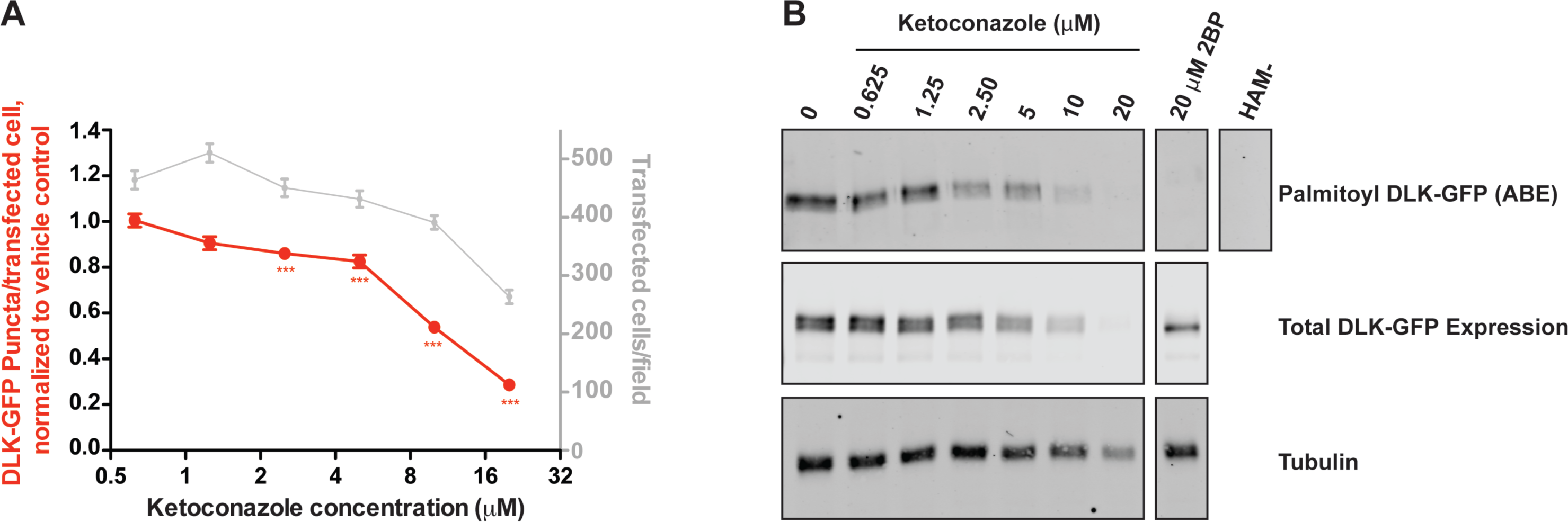
Dose-dependent inhibition of DLK-GFP localization and palmitoylation by ketoconazole. **A**) Quantified DLK puncta/per transfected cell from HEK293T cells transfected as in Fig 2 and treated with the indicated concentrations of ketoconazole. Data are plotted relative to DMSO vehicle control. **B**) HEK293T cells in 6 cm plates were transfected with DLK-GFP cDNA and subsequently treated with the indicated concentrations of ketoconazole. Palmitoyl-proteins were purified from lysates using Acyl Biotin Exchange. Total levels of DLK and tubulin were detected by western blotting of parent lysates. A negative control ABE sample processed in the absence of the key reagent hydroxylamine (HAM-) was generated by combining equal fractions of lysates from all conditions.

To determine whether effects of ketoconazole on DLK localization were linked to reduced palmitoylation, we subjected lysates of DLK-GFP-expressing cells to an orthogonal mechanism of action assay, Acyl-Biotin Exchange (ABE). In this assay, thioester-linked acyl modifications (i.e. palmitoylation) are exchanged for biotin and the resultant biotinyl-proteins are affinity-purified from cell lysates using avidin-conjugated beads [11, 22]. Consistent with our findings from the DLK-GFP localization assay, ketoconazole dose-dependently decreased palmitoylation of DLK-GFP in ABE assays (Figure 3B). At 2.5 μM ketoconazole predominantly affected DLK palmitoylation, while at 5 μM and 10 μM, ketoconazole reduced DLK palmitoylation to a greater extent but also slightly reduced total levels of DLK-GFP expression. At >10 μM, ketoconazole reduced tubulin levels, consistent with the reduced mCherry-NLS counts seen in the DLK-GFP localization assay, and again suggesting possible effects on transcription/translation and/or cytotoxicity. However, findings from the ABE assay broadly mirrored those from our primary assay, with results from both assays suggesting that there is a window within which ketoconazole reduces both DLK punctate localization and palmitoylation without broadly affecting protein transcription, translation or stability.

### Ketoconazole inhibits palmitoylation of DLK and PSD-95, but not GAP43

We next sought to assess whether ketoconazole specifically reduces DLK palmitoylation levels, or affects cellular palmitoylation more broadly. To address this question, we used ABE to assess palmitoylation of two other well characterized palmitoyl-proteins, Growth-Associated Protein-43 (GAP-43) and Post-Synaptic Density-95 (PSD-95) [23, 24]. In transfected HEK293T cells, ketoconazole significantly decreased palmitoylation of both DLK-GFP and PSD-95 (Figure 4A, C), but did not reduce GAP43-Myc palmitoylation (Figure 4B). In parallel assays, the broad spectrum palmitoylation inhibitor 2BP reduced palmitoylation of all three proteins. These findings suggest that ketoconazole is not a broad spectrum inhibitor of protein palmitoylation and is thus distinct from 2BP.

**Figure 4.**
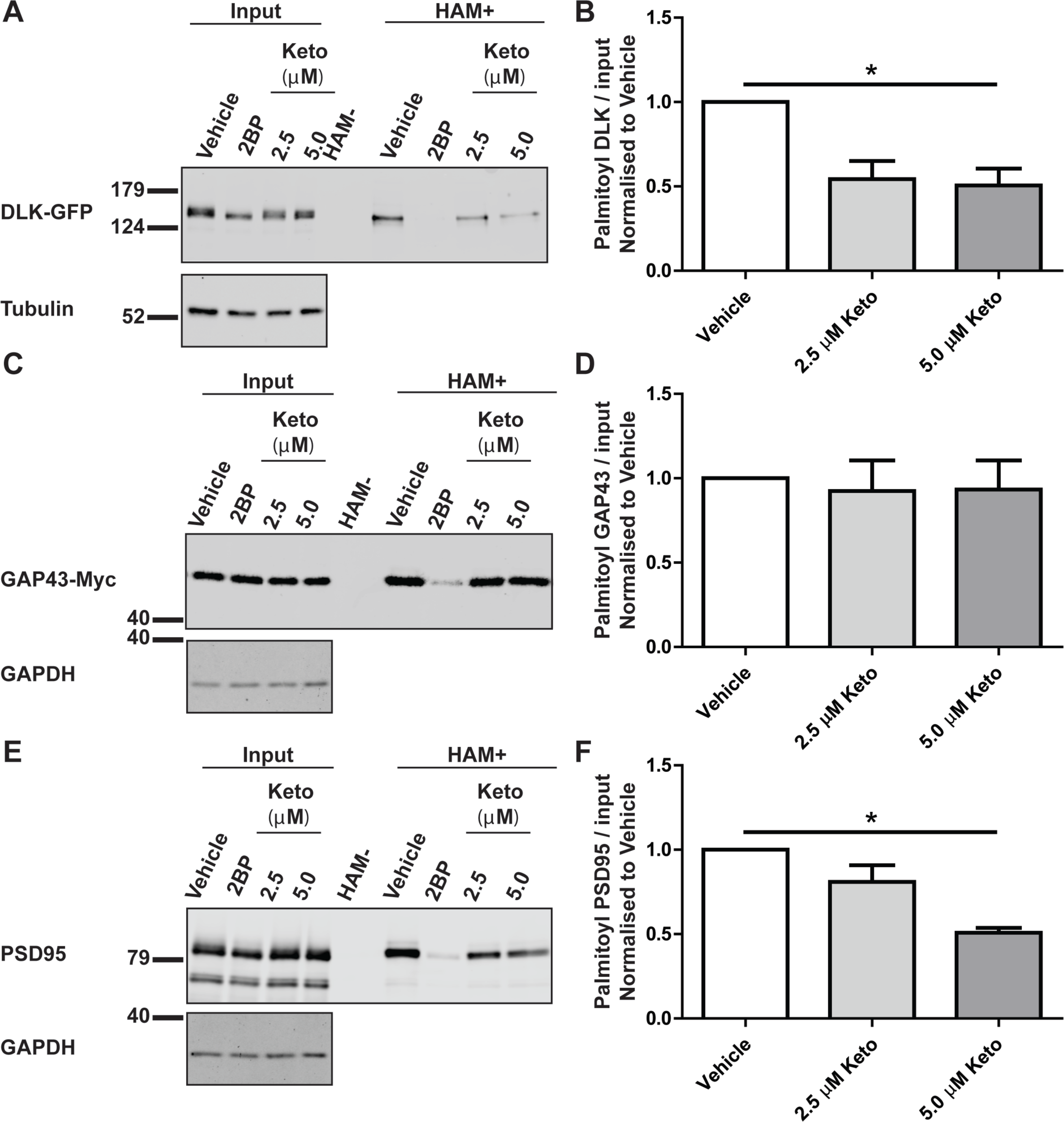
Ketoconazole inhibits palmitoylation of DLK and PSD-95, but not GAP43. **A)** HEK293T cells were transfected with DLK-GFP cDNA and incubated with 20 μM 2BP, or with 2.5 μM or 5 μM ketoconazole 2 h post-transfection for 16–18 h. Upper western blot shows DLK total expression and palmitoyl-DLK levels (from ABE, ‘HAM+’) for each condition. A control ABE sample processed in the absence of the key reagent hydroxylamine (HAM-) was generated by combining equal fractions of lysates from all conditions. Lower western blot shows tubulin levels, an indication of total protein expression. **B**) Histogram of pooled data (mean +SEM) for 4 determinations per condition from *A*. Ketoconazole and 2BP both significantly reduce DLK palmitoylation. **C**) As A, except that cells were transfected with GAP43-Myc cDNA ABE fractions were blotted with anti-Myc antibody and cell lysates were blotted to detect total expression of GAP-43-myc (upper panel) and GAPDH (lower panel). **D**) Histogram of pooled data (mean ±SEM) for 4 determinations per condition from *C*. Ketoconazole does not reduce GAP-43 palmitoylation, but 2BP does. **E**) As C, except that cells were transfected with PSD-95 cDNA and total lysates and ABE fractions were blotted with anti-PSD-95 antibody. **F**) Histogram of pooled data (mean ±SEM) for 4 determinations per condition from *E*. 5 μM ketoconazole and 2BP both reduce PSD-95 palmitoylation. One-way ANOVA, Kruskal-Wallis post-hoc analysis; **A**) ANOVA p=0.0214, h=7.692, **C**) ANOVA not significant, **E**) ANOVA p=0.0158, h=8.290.

### Ketoconazole inhibits DLK-dependent cJun phosphorylation in sensory neurons subjected to trophic deprivation

Given the importance of palmitoylation for DLK-dependent signalling [11] and the clear effects of ketoconazole on DLK palmitoylation levels (Figure 3), we next assessed whether ketoconazole could affect neuronal DLK signalling. Sensory neurons subjected to Trophic Deprivation (TD) activate a pro-degenerative DLK-JNK signaling pathway that leads to the phosphorylation of the transcription factor c-Jun [25, 26]. Consistent with our prior finding that c-Jun phosphorylation requires palmitoyl-DLK [11], 2BP completely prevented TD-induced c-Jun phosphorylation (Fig. 5). Interestingly, ketoconazole also significantly reduced TD-induced c-Jun phosphorylation in sister cultures subjected to TD (Fig. 5). These findings suggest that ketoconazole can reduce not only DLK localization and palmitoylation, but also DLK-dependent neuronal signalling.

**Figure 5.**
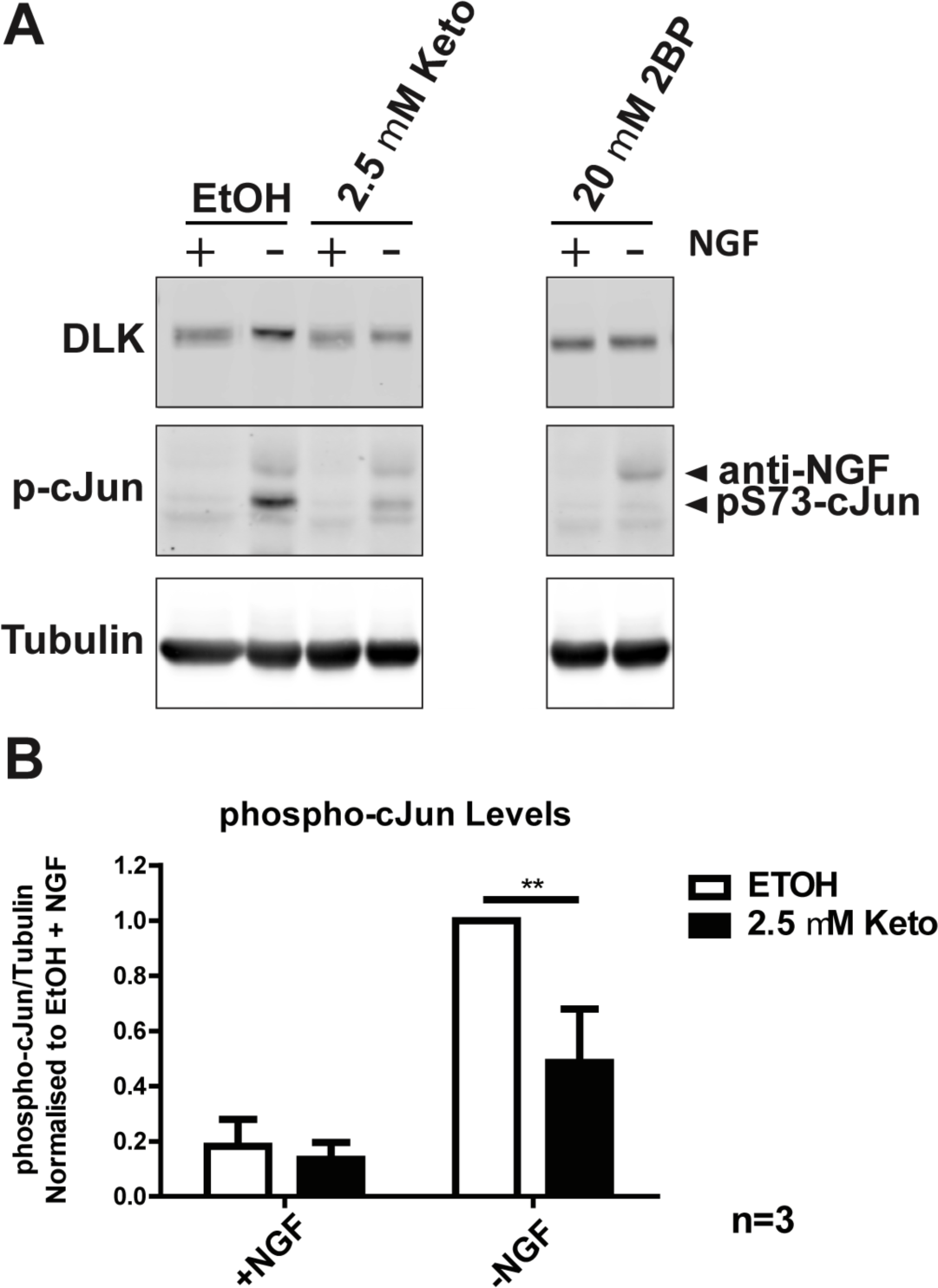
Ketoconazole significantly decreases DLK-mediated phospho-cJun activation in primary neurons. **A**) Dorsal Root Ganglion (DRG) neurons were treated at 7 Days *in vitro* (DIV 7) with 2.5 μM Ketoconazole overnight or 20 μM 2BP for 2 h prior to a 2.5 h NGF withdrawal in presence of the indicated compound. Cells were lysed in SDS-PAGE loading buffer and levels of endogenous DLK, phospho-cJun and tubulin were detected by western blot. **B**) Quantification of phospho-cJun normalised to –NGF vehicle treated cells. Two-way ANOVA indicates significant effects of interaction (p=0.0071), NGF (p=0.0026) and Ketoconazole (p=0.0001). The effect of Ketoconazole in DRGs undergoing NGF withdrawal was also significant as determined by the Bonferroni post-test (p<0.01). Error bars represent SEM.

## Discussion

There is considerable interest in inhibiting DLK signaling as a therapeutic strategy to prevent neurodegeneration in a variety of pathological conditions. Although direct inhibitors of DLK’s kinase activity are being developed [1, 8–10], a complementary neuroprotective approach might be to prevent DLK palmitoylation, because this lipid modification is essential for DLK’s kinase activity [11]. Our high content imaging screen facilitates this latter approach by exploiting dramatic palmitoylation-dependent changes in DLK localization to identify compounds that inhibit DLK palmitoylation. Our screening method is robust and is capable of identifying compounds that reduce DLK palmitoylation in orthogonal biochemical assays and which also reduce DLK-dependent pro-degenerative signaling in neurons. Moreover, because inhibition of palmitoylation has not been pursued as a neuroprotective strategy, our screening platform has the potential to identify novel classes of compounds that may have considerable therapeutic potential.

Our initial screen and follow-up assays represent an important proof of principle, but several questions remain to be addressed. In particular, although our top ‘hit’ ketoconazole markedly affected DLK localization in our primary screen, how this compound acts to reduce DLK palmitoylation and signaling remains to be determined. Nonetheless, findings from some of our additional experiments can help rule out certain possible explanations as to ketoconazole’s mechanism of action.

For example, ketoconazole reduces palmitoylation of DLK and, to a lesser extent, PSD-95, but does not affect palmitoylation of GAP-43. These findings stand in contrast to the broad spectrum palmitoylation inhibitor 2BP, suggesting that ketoconazole and 2BP act *via* different mechanisms.

Our results also provide insights into ketoconazole’s mechanistic specificity. For example, ketoconazole reduces palmitoylation of DLK, which is palmitoylated at a single, internal site, and of PSD-95, which undergoes N-terminal dual palmitoylation. It thus seems unlikely that ketoconazole’s action is defined by the number or location of palmitoylation sites in a given palmitoyl-protein. We speculate that ketoconazole may instead inhibit one or more PAT(s) that can palmitoylate DLK and PSD-95. However, both PSD-95 and DLK can be palmitoylated by a number of different PATs in transfected cells [11, 27], and HEK293T cells express all 23 human PATs [28, 29], so testing this possibility is far from trivial.

It is also informative to consider prior descriptions of ketoconazole’s activity in other contexts. Ketoconazole is an antifungal that was first identified as an inhibitor of enzymes involved in generating ergosterol, the fungal form of cholesterol [30]. In humans, ketoconazole inhibits testicular androgen production and can inhibit the 17α-hydroxylase and 17,20-lyase activities of the steroidogenic P450 enzyme Cytochrome P450 17 A1 (CYP17A1 [31]). However, how these activities relate to ketoconazole’s effects on DLK palmitoylation and signaling is unclear.

The chemical moiety within ketoconazole that acts to reduce palmitoylation is also not fully clear. Interestingly, a second azole-containing compound, sulconazole, was also identified in our screen (Table 1). However, miconazole, a third azole-containing compound that is included in the Prestwick library, was not identified in the screen and was ineffective at reducing DLK puncta at the same concentration as ketoconazole and was toxic at higher concentrations in follow-up assays (D.D.O.M., unpublished observations). Further chemi-informatic analysis may facilitate identification of possible common functional moieties present in ketoconazole and/or other screen hits.

While our assay is designed to identify compounds that prevent DLK palmitoylation, it can also be used to identify compounds that specifically reduce DLK stability. Indeed, ketoconazole’s ability to reduce numbers of DLK puncta in our initial screen may in part be due to this secondary activity, because at 10 μM (the concentration used in the initial screen) ketoconazole did slightly reduce total protein expression of DLK-GFP. However, this additional capability of the screening platform is actually an unexpected bonus - given DLK’s role as a key controller of neurodegeneration, compounds that act to destabilize DLK and/or increase DLK degradation might also be of considerable therapeutic benefit. The tight control of DLK levels by ubiquitin-dependent degradation [32–34] suggests that our screen could also identify activators of DLK ubiquitylation and/or inhibitors of DLK de-ubiquitylation. Compounds that act in either of these ways would also be of considerable interest therapeutically.

Finally, dramatic palmitoylation-dependent changes in protein subcellular localization are well described not just for DLK but key regulators of axon integrity, synaptic transmission / higher brain function, and cell growth / proliferation [23, 35–40]. Fluorescent- or other epitope-tagged versions of many of these proteins are either readily available or can be easily generated, making HCS a powerful approach to identify small molecules that could affect their localization and activity. While we focused on palmitoylation-dependent changes in the number of DLK puncta per cell, other aspects of protein subcellular localization that can be controlled by palmitoylation (e.g. plasma membrane targeting) can also be quantified by ImageXpress, or by similar software [41]. Our screening platform could thus be readily adapted to identify compounds that affect the palmitoylation-dependent targeting of a variety of therapeutically important proteins (e.g. Ras, oncogenic Src family kinases [40, 42, 43]) to other subcellular locations. In addition, HCS is also readily compatible with genome-wide RNAi or CRISPR-based screening [17, 44], so our screening platform could be combined with these methods to identify upstream regulators (e.g. PATs and/or thioesterases, or specific binding partners) that control palmitoyl-protein subcellular localization. Such approaches have considerable potential both to provide new biological insights into the control of protein palmitoylation, and also to identify compounds and therapeutic targets to lessen the impact of numerous pathological conditions.

## Materials and Methods

The following antibodies, from the indicated sources, were used in this study: Rabbit anti-GFP (Invitrogen / Thermo Fisher Biosciences: catalog #A11122); mouse anti-GM130 (BD Biosciences, Catalog #610822); Rabbit anti-phospho c-Jun Ser-73 (Cell Signaling Technology, Catalog #3270); DLK/MAP3K12 (Sigma/ Prestige, Catalog #HPA039936); mouse anti-PSD-95 (Antibodies Inc., Catalog #75-028); Myc 9E10 (University of Pennsylvania Cell Center, Catalog #3207), rabbit anti-GAPDH (Thermo Scientific, Catalog #PA1-987), mouse anti-tubulin (Millipore Sigma, Catalog #T7451), sheep anti-NGF (CedarLane, catalog #CLMCNET-031). Wild type DLK-GFP and palmitoyl mutant (C127S) DLK were previously described [11]. pmCherry-NLS was a gift from Martin Offterdinger (Addgene Plasmid #39319) [20]. Rat GAP43 cDNA was gene synthesized (Genewiz) and subcloned into the vector FEW [11] upstream of a C-terminal myc tag. Untagged PSD-95 cDNA was a gift from Dr. R.L. Huganir (Johns Hopkins University Medical School) [45]. 2-Bromopalmitate (2BP) and S-Methyl methanethiosulfonate (MMTS) were from Sigma. The Prestwick Chemical library was purchased by Temple University’s Moulder Center for Drug Discovery and formulated as 10 mM stocks in DMSO. Fresh Ketoconazole stock for follow-up assays was from LKT Laboratories (Catalog #K1676). All other chemicals were from Fisher Biosciences and were of the highest reagent grade.

### Cell transfection

HEK293T cells were transfected using a calcium phosphate-based method as described previously [36].

### Transfection and fixation of cells for microscopy

In initial experiments of DLK-GFP subcellular localization, HEK293T cells seeded on poly-lysine-coated coverslips (Warner Instruments) in 6 cm dishes were transfected as above. Cells were treated with 100 μM 2BP 5h later and then fixed 8h post-transfection in 4% (wt/vol) paraformaldehyde, 4% (wt/vol) sucrose in PBS. After PBS washes, cells were permeabilized with PBS containing 0.25% (wt/vol) Triton X-100, blocked in 10% (vol/vol) normal goat serum (NGS) in PBS and incubated overnight at 4°C with rabbit anti-GFP and mouse anti-GM130 antibodies in 10% (vol/vol) NGS, followed by incubation with AlexaFluor-conjugated secondary antibodies for 1 h at room temperature. Nuclei were stained with 300 nM DAPI in PBS for 10 min and coverslips were mounted in Fluoromount G (Southern Biotech) before imaging.

For High Content Screening assays, HEK293T cells were plated in poly-lysine coated 96 well plates (Greiner Bio-One, black walled chimney-wells), transfected as above and treated with 2BP (10 μM final concentration), library compounds or DMSO vehicle control at 2h post-transfection. The Prestwick Compound Library was spotted onto 96 well plates at 10 mM in DMSO and resuspended in 200 μL DMEM. 40 μL of diluted drug was then added to cells in 160 μL of DMEM (containing glutamax, 10% FBS and antibiotics) in duplicate. Cells were incubated with compounds at 37°C for a further 14 h. Subsequently, medium was removed and cells fixed in 4% PFA (1x PBS) for 20 mins at RT, washed twice with PBS and stained with 300 nM DAPI for 30 mins at RT, followed by 2 washes of PBS.

### High Content Screening

High Content screening was performed using the ImageXpress micro high content imaging system (Molecular Devices, Downingtown, PA) driven by MetaXpress software. Six images per well were acquired in each of three channels (DAPI, FITC, TRITC) at 10X magnification in an unbiased fashion. Images were analyzed using the MetaXpress ‘Multiwavelength Scoring’ (for mCherry-NLS signals) and ‘Transfluor’ modules (for DLK-GFP signals). Data were exported to Excel utilizing the AcuityXpress software package (Molecular Devices).

### Thresholding

Compounds that reduced either DAPI or NLS signals by greater than 30% of the average of vehicle-treated controls for each day were excluded from analysis due to likely cytotoxicity and/or broad effects on transcription, translation or protein stability. MetaXpress imaging software was then used to determine the effect of the remaining compounds on DLK puncta (“Total Puncta Count” option, from DLK-GFP signal) and total number of transfected cells (from mCherry-NLS signal). The term “DLK puncta per transfected cell” was used for this readout because punctate DLK-GFP distribution is likely a mixture of Golgi-associated and vesicle-associated pools of DLK. Compounds that reduced DLK puncta per transfected cell by 3 times the standard deviation of the mean of all vehicle-treated controls were considered “Hits”.

### Follow-up Assay

To confirm the effect of ketoconazole, HEK293T cells were seeded on poly-lysine-coated 96 well plates, transfected as above with DLK-GFP and mCherry-NLS cDNAs. Two hours post-transfection, cells were treated with freshly dissolved ketoconazole over a 5-point dilution range, or with DMSO vehicle control. Cells were fixed 16h later and processed and imaged as for the primary screen.

### Palmitoylation assay

Palmitoylation of transfected proteins in HEK293T cells was assessed by acyl biotin exchange assays, as previously described [36] except that bands were imaged and quantified using a LiCOR Odyssey system. Images were prepared and analyzed using Image Studio Lite Ver 4.0.

### NGF Withdrawal

Primary dorsal root ganglion (DRG) were prepared from embryonic day 14.5 rat embryos, as previously described [11]. At 7 days *in vitro* DRGs were pretreated with 2.5 μM Ketoconazole overnight or 20 μM 2BP for 2 h prior to withdrawal of NGF in the presence of NGF antibody in the continued presence of drug. Cells were then lysed in SDS-PAGE loading buffer and processed for subsequent SDS-PAGE and subsequent immunoblotting. Images were acquired and analyzed as above.

### Statistical Analysis

Where indicated, the non-parametric one-way ANOVA Kruskal-Wallis test was performed with a Dunn’s multiple comparison *post-hoc* analysis. In addition, 2-way ANOVA was performed with Bonferroni *post-hoc* analysis. All error bars represent SEM.

## Acknowledgements

The authors thank Drs. Jingwen Niu and Francesca DeSimone for help with neuronal cultures, and for preparing GAP43-Myc, respectively, and John Gordon for assistance with screening compounds. This work was supported by Shriners Hospital for Children Grant #87400 PHI and NIH Grant #R01 NS094402 (both to G.M.T.). S.S.S is a Brody Family Medical Trust Fund Fellow.

---

**Supplementary Figure 1. Additional readouts of DLK punctate distribution are also highly palmitoylation-dependent**. Images of DLK-GFP expressing HEK293T cells from Figure 1C were analyzed using the ‘Transfluor’ modules within MetaXpress analysis software to quantify changes in different aspects of punctate DLK signals, in particular “Puncta Count,” “Puncta Count Per Cell,” “Puncta Total Area,” “Puncta Area Per Cell,” “Puncta Integrated Intensity,” and “Puncta Average Intensity”, as indicated. Error bars represent SD. Z-prime value (z′) for each metric is indicated. All measurements reached a significance of 0.0001 by unpaired parametric two-tailed t-test.

